# Liquid-crystalline lipid phase transitions in lipid droplets selectively remodel the LD proteome

**DOI:** 10.1101/2021.08.30.458229

**Authors:** Sean Rogers, Long Gui, Anastasiia Kovalenko, Evan Reetz, Daniela Nicastro, W. Mike Henne

**Author notes:** these authors contributed equally to this work (Sean Rogers, Long Gui, Anastasiia Kovalenko).

## Abstract

Lipid droplets (LDs) are reservoirs for triglycerides (TGs) and sterol-esters (SEs). How lipids are organized within LDs and influence the LD proteome remains unclear. Using *in situ* cryo-electron tomography, we show that glucose restriction triggers lipid phase transitions within LDs generating liquid-crystalline lattices inside them. Mechanistically, this requires TG lipolysis, which alters LD neutral lipid composition and promotes SE transition to a liquid-crystalline phase. Fluorescence imaging and proteomics further reveal that LD liquid-crystalline lattices selectively remodel the LD proteome. Some canonical LD proteins including Erg6 re-localize to the ER network, whereas others remain on LDs. Model peptide LiveDrop also redistributes from LDs to the ER, suggesting liquid-crystalline-phases influence ER-LD inter-organelle transport. Proteomics also indicates glucose restriction elevates peroxisome lipid oxidation, suggesting TG mobilization provides fatty acids for cellular energetics. This suggests glucose restriction drives TG mobilization, which alters the phase properties of LD lipids and selectively remodels the LD proteome.

## Introduction

Lipid droplets (LDs) are unique endoplasmic reticulum (ER)-derived organelles dedicated to the storage of energy-rich neutral lipids. Structurally LDs are composed of a hydrophobic core of triglycerides (TGs) and sterol-esters (SEs) that is surrounded by a phospholipid monolayer that either contains or is decorated by specific proteins. Beyond their roles in energy homeostasis, recent work highlights the roles of LDs in signaling, development, and metabolism (Welte and Gould, 2017), (Olzmann and Carvalho, 2019), (Walther et al., 2017). These diverse jobs are largely dictated by the LD proteome, but a pervasive question is how specific proteins are targeted to the LD surface. Furthermore, whether the LD proteome is static or dynamic, and how metabolic cues influence LD protein residency is poorly understood.

LDs are generated at the ER and often remain connected to the ER bilayer for extended periods (Jacquier et al., 2011), (Kassan et al., 2013). As such, Type I LD proteins can translocate between the ER and LD monolayer via lipidic bridges connecting the two organelles (Wilfling et al., 2013). Elegant *in vitro* studies have suggested that LD localization promotes energetically favorable conformational changes within some proteins, and the movement of proteins to LDs from the ER network can even influence their enzymatic activities, or modulate their degradation (Caillon et al., 2020), (Chorlay and Thiam, 2020), (Leber et al., 1998), (Schmidt et al., 2013), (Ohsaki et al., 2006). A second mechanism of LD targeting occurs from the cytoplasm, where soluble proteins insert into the LD monolayer via a hydrophobic region, amphipathic helix, or lipid moiety. Here hydrophobic protein regions recognize packing defects between the phospholipid monolayer lipid head groups, enabling their insertion into the neutral lipid core (Chorlay and Thiam, 2020).

Although monolayer phospholipids can regulate LD protein targeting, how *neutral lipids* influence protein localization is less understood. However, neutral lipids clearly impact the composition of the LD surface proteome; for example, in yeast, some proteins preferentially decorate TG-rich LDs (Gao et al., 2017). Molecular studies also indicate that protein insertion into the LD neutral lipid core enables proteins to fold with lower free energy, and polar residues within hydrophobic regions can even interact with TG, further anchoring them to the LD (Olarte et al., 2020). However, how neutral lipid pools ultimately influence the composition and dynamics of the LD proteome is relatively unexplored, yet central to our understanding of LD organization and functional diversity.

Neutral lipids generally form an amorphous mixture within the hydrophobic LD core. This organization can change in response to various cellular stimuli. HeLa cells induced into mitotic arrest or starvation exhibit lipid phase transitions within their LDs, generating liquid-crystalline lattices (LCLs) inside LDs with a striking onion-like appearance by cryo-electron tomography (cryo-ET) (Mahamid et al., 2019). Yeast biochemical studies also proposed similar segregation of TGs and SEs into discrete layers within LDs (Czabany et al., 2008). This lipid reorganization is attributed to the biophysical properties of SEs, which can transition from disordered to ordered smectic phases under physiological conditions (Kroon, 1981), (Ginsburg et al., 1984), (Shimobayashi S, 2019), (Czabany et al., 2008). Such phase transitions are also associated with human pathologies including atherosclerosis, and liquid-crystalline LDs were even observed in the macrophage of a patient with Tangier disease (Lundberg, 1985), (Katz et al., 1977). How these phase transitions are triggered, however, and whether they influence organelle physiology, or are simply a biophysical consequence of the properties of SEs, is unknown.

Here, we utilized budding yeast to dissect the metabolic cues governing lipid phase transitions within LDs. We used cryo-ET of cryo-focused ion beam (cryo-FIB) milled yeast cells to study the *in situ* architecture of LDs in their native environment, under ambient or glucose-starved conditions. We show that in response to acute glucose restriction, yeast initiate TG lipolysis, which induces the formation of LCLs within LDs. In line with this lipid mobilization, global proteomics reveals that glucose restriction promotes metabolic remodeling favoring peroxisome fatty acid oxidation and mitochondrial metabolism. Furthermore, we find LD liquid-crystalline remodeling selectively changes the LD surface proteome, promoting the redistribution of some proteins from the LD surface to the ER network while others are retained on LCL-LDs.

## Results

### Acute glucose restriction promotes TG lipolysis-dependent liquid-crystalline phase transitions in LDs

Previous studies from our group indicated that budding yeast exposed to acute glucose restriction (AGR), where yeast are transferred from a glucose-rich (2%) synthetic complete media to a low-glucose (0.001%) media, exhibit metabolic remodeling that favors the production of SEs, which are stored in LDs (Rogers et al., 2021). We used cryo-ET to investigate if AGR also impacts LD morphology. We rapidly froze yeast cells that were either in logarithmic (log-phase) growth in glucose-rich media, or exposed to 4 hrs of AGR, and used cryo-FIB milling to generate 100-200-nm-thick lamellae of the vitrified cells. These lamellae were then imaged by cryo-ET to reveal the three-dimensional (3D) structure of native LDs *in situ*. The cryo-FIB milled lamella exhibited a well-preserved yeast ultrastructure, including the nucleus, vacuole, mitochondria and LDs (**Figure 1A, SFigure1, SMovie 1**). Typical LDs could be distinguished from other cellular organelles by their relatively electron-dense, amorphous interior that was surrounded by a thin phospholipid monolayer (**Figure 1B, SMovie 2**). In contrast to normal LDs in glucose-fed log-phase cells, ~77% of the LDs observed in 4hrs AGR-treated yeast displayed reorganization of their interior, including the appearance of distinct concentric rings in the LD periphery (**Figure 1 C-D, M for quantification, SMovie 3**). These rings appear similar to lattices previously observed in liquid-crystalline-phase LDs, which exhibited a regular spacing of ~3.4-3.6nm between their layers, suggesting they were composed of sterol-esters (Mahamid et al., 2019), (Engelman and Hillman, 1976). Indeed, our line-scan analysis showed a regular 3.4nm spacing between rings (**Figure 1E**), suggesting these LDs exhibited liquid-crystalline lattices (LCLs). Thus, we refer these “onion-like” LDs as LCL-LDs. Notably, these were never observed in the log-phase yeast (**Figure 1B, M**).

**Figure 1:**
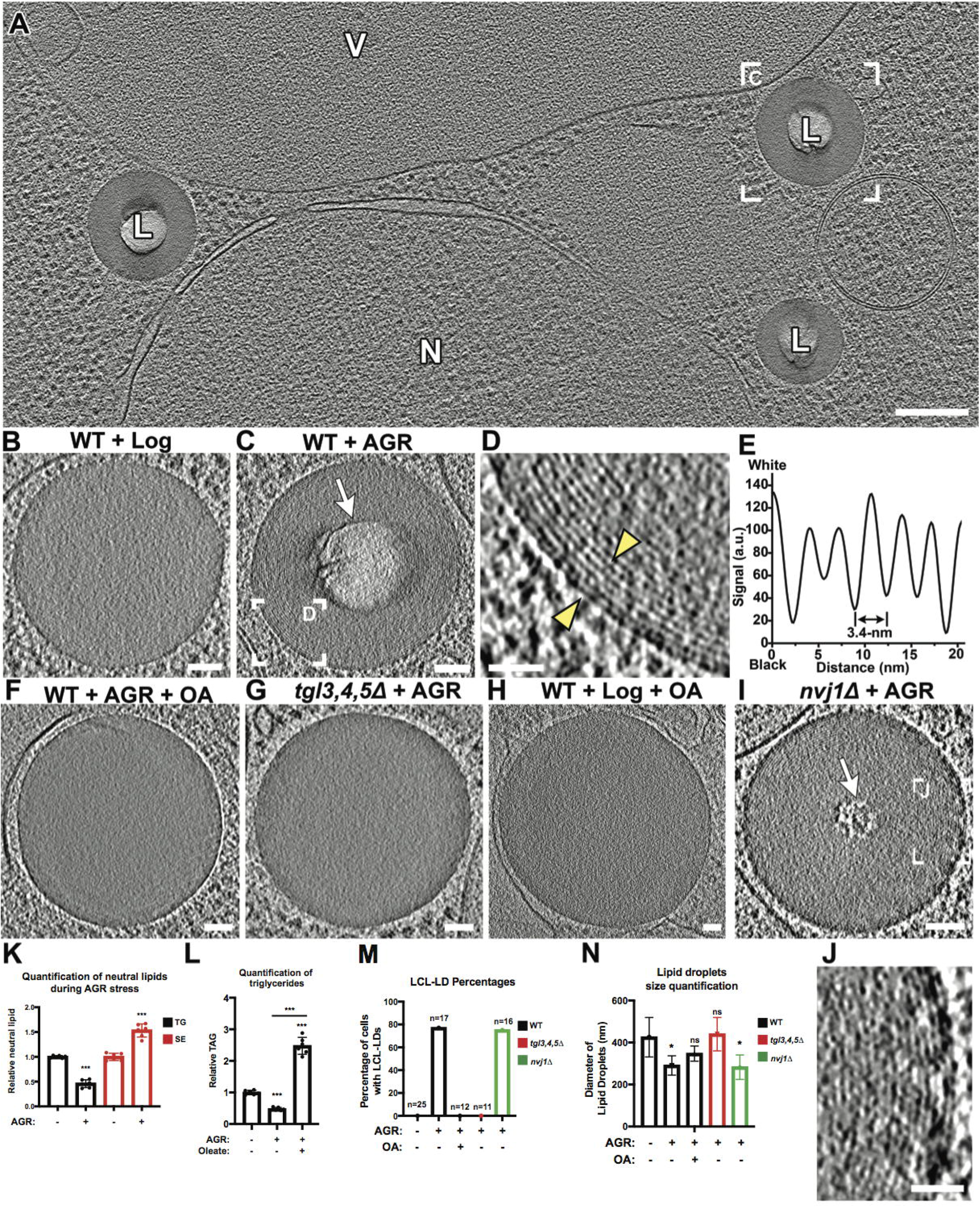
Visualization of the liquid-crystalline layers in lipid droplets (LCL-LD) promoted by TG lipolysis using *in situ* cryo-ET. **A**) Representative tomographic slice from a cryo-FIB-milled and cryo-ET reconstructed wildtype (WT) yeast cell grown for 4hrs under acute glucose restriction (AGR). Note the “bubbled” (lighter) centers of the LDs (L). V, vacuole. N, nucleus. A different tomographic slice of the boxed LD is also shown in (**C**). **B-J**) Representative tomographic slices of LDs in yeast from glucose-fed WT in log phase (**B**), WT after 4hrs AGR (**C**, boxed area magnified in **D, E** shows line-scan plot of area between yellow arrowheads), WT after 4hrs AGR + 0.1% oleate (OA) (**F**), *tgl3,4,5Δ* yeast after 4hrs AGR (**G**), WT cultured with 2% glucose and 0.1% OA (**H**), *nvj1Δ* after 4hrs AGR (**I**, and boxed area magnified in **J**). Liquid-crystalline layers (LCL) were only observed in LDs from WT and *nvj1Δ* yeasts in AGR (**C**, **D**, **I**, **J**). White arrows highlight the ‘bubbles’ due to electron radiation in centers of LCL-LDs. **K**) Quantification of relative whole-cell TGs and SEs in log and 4hrs AGR conditions. **L**) Relative TGs in log and 4hrs AGR conditions. **M, N**) % abundance of LCL-LDs (**M**) and diameters of LDs (**N**) under various conditions measured in cryo-tomograms. Note that the observed diameter depends on the plane at which the LDs were sectioned; therefore, for size measurements, only LDs with clearly visible monolayer (indicating a slice through the LD center) were included. Scale bars: 200nm (A). 50nm (B-C, F-I). 20nm (D, J).

In addition to the peripheral lattices, the amorphous center of LCL-LDs was unusually sensitive to electron radiation, causing excessive radiolysis and “bubbling” (i.e. the generation of a gas bubble trapped in the ice that appears white in cryo-EM images) during tilt-series acquisition (**Figure 1C, white arrow**). This increased radiation sensitivity was only observed in LCL-LDs, but not in LDs with entirely amorphous lumen (i.e. not observed in the 23% unordered LDs of AGR-treated yeast, nor in any LDs of log-phase yeast). We generated comparative ‘bubblegrams’, (i.e. a series of 2D cryo-EM images where the same sample area was exposed to an increasing amount of electron dose), which revealed that the centers of LCL-LDs exhibited bubbling following exposure to <30 e/Å^2^, whereas amorphous LDs from log-phase yeast did not show any bubbling even at 400 e/Å^2^ dosages (**SFigure1 A-J**). Previous studies of electron radiation-induced bubbling of frozen biomolecules in aqueous solution and cells demonstrated that similar gas bubbles contained mostly molecular hydrogen gas (Leapman and Sun, 1995) (Aronova et al., 2011). Although the mechanism of radiation-induced bubbling and increased radiation-sensitivity within the center of LCL-LDs is not clear, it may be due to the production of gases derived from a specific combination of lipids or metabolites present within LCL-LDs.

To investigate the effects of AGR stress on yeast neutral lipid pools, we monitored TG and SE levels in log-phase and 4hrs AGR-treated yeast. Indeed, AGR treated yeast contained significantly less TGs (**Figure 1K**). As expected, AGR yeast also had increased amounts of SEs (**Figure 1K)**, as previously observed (Rogers et al., 2021), indicating the TG:SE ratio within the LDs was significantly decreased to ~0.5:1.5 compared to a normal ratio of ~1:1 (Leber et al., 1994). We hypothesized that LCL-LD formation was promoted by TG loss from LDs. To test this, cryo-ET was performed on yeast lacking the major TG lipases (*tgl3,4,5Δ*). Indeed, 4hrs AGR treated *tgl3,4,5Δ* yeast did not form any detectable LCL-LDs (**Figure 1G, M**), suggesting TG lipolysis was required for LCL-LD formation. In support of this, LDs in wildtype (WT) AGR-treated yeast were significantly smaller in diameter than log-phase LDs, and this reduced size was suppressed in *tgl3,4,5Δ* yeast (**Figure 1N**), suggesting the size reduction was due to lipid loss via TG lipolysis.

To further dissect how TGs influence LCL-LDs, we treated yeast with 0.1% oleic acid (OA), which promotes TG synthesis. As expected, OA elevated cellular TG levels in yeast when they were cultured in it during 4hrs AGR treatment (**Figure 1L**), and notably no LCL-LDs were observed during log-phase nor in this AGR condition (**Figure 1F, H, M**). In line with this, whereas LD sizes in AGR-treated yeast were significantly smaller than in log-phase cells, their sizes slightly recovered under the AGR plus OA condition (**Figure 1F, N**). Since we previously observed that the nucleus-vacuole junction (NVJ) can serve as a site for LD biogenesis during nutrient stress (Hariri et al., 2018), we also examined whether NVJ loss impacted LCL-LD formation. Cryo-ET of *nvj1Δ* yeast cells showed the expected loss of tight contacts between the outer nuclear envelope and the vacuole (**SFigure 1K, L**). However, *nvj1Δ* yeast exhibited ~75% LCL-LDs under AGR conditions, indicating that the NVJ was not required for LCL-LD formation (**Figure 1I, J, M**).

Since SEs can form liquid-crystalline lattices, we tested whether SEs were required for LCL-LD formation. We monitored LDs in *are1are2Δ* yeast that cannot synthesize SEs. Surprisingly, in 15 different cryo-FIB lamella of *are1are2Δ* yeast cells no LDs could be observed (**SFigure 1M**). However, fluorescence staining with monodansylpentane (MDH) LD stain confirmed the presence of LDs in *are1are2Δ* yeast during AGR stress, but they were small and sparse in many yeast compared to any of the other examined strains (**SFigure 1N**). The reduction in LD size and abundance may account for the inability to observe LDs in the cryo-tomograms of the 100-200nm thick lamellae.

Collectively, these data suggest that TG abundance is a key modulator of the SE phase transitions within the LD, and indicate Tgl-dependent TG lipolysis during AGR promotes LCL-LD formation by depleting the TG pool that maintains SE in its disordered phase.

### LCL-LD formation selectively remodels the LD proteome

While studies indicate that LD proteins may interact with TGs contained within the LD interior (Olarte et al., 2020), (Santinho et al., 2021), it is unknown whether smectic lipid phase transitions influence LD protein targeting. Therefore, we imaged the canonical LD protein Erg6 tagged with mNeonGreen (Erg6-mNg) over time in AGR conditions. As expected, Erg6-mNg initially colocalized with LD stain at the start of AGR (t=0). However, the Erg6 labeling pattern changed after ~1hr AGR, and primarily decorated the cortical ER and nuclear envelope (**Figure 2A**). Erg6-mNg remained at the ER network throughout 2, 4, and 24 hrs AGR, and notably the LD stain gradually dimmed over these time-points, consistent with the loss of LD volume via lipolysis. Remarkably, the addition of 0.1% OA, or genetic ablation of TG lipases both rescued Erg6-mNg LD targeting at 4hrs AGR (**Figure 2B, SFigure 2A**). Since our cryo-ET results showed lack of LCL-LD formation in these conditions, it suggested that Erg6-mNg de-localization from LDs tightly correlates with LCL-LD formation.

**Figure 2:**
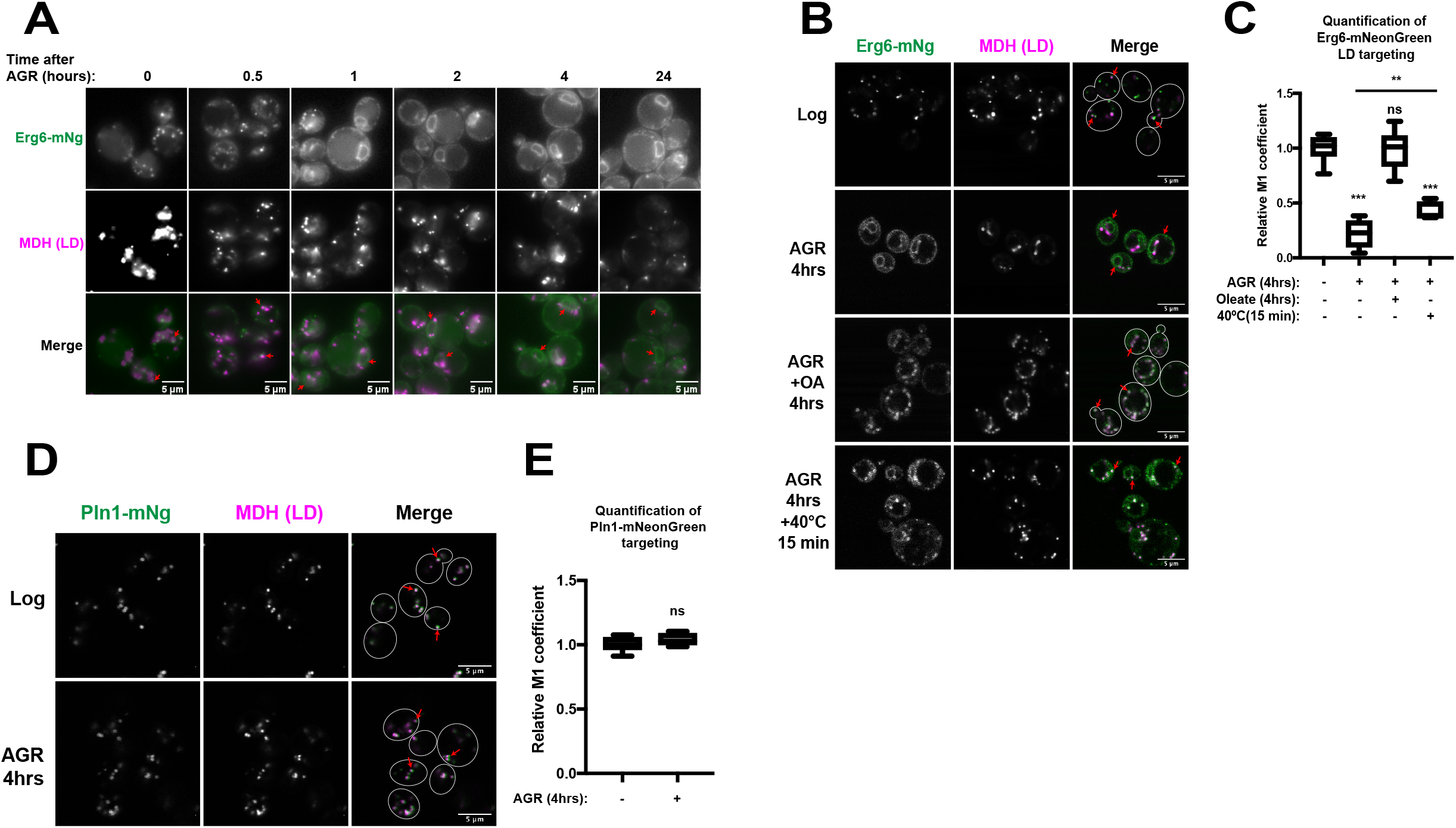
Erg6 LD de-localization correlates with LCL-LD formation. **A**) Yeast expressing Erg6-mNeonGreen (mNg) and stained for LDs (monodansylpentane, MDH) at time-points when yeasts were transferred from log-phase (2% glucose) to acute glucose restriction (AGR). Red arrows indicate protein targeting. **B**) Yeast with Erg6-mNg and LD/MDH stain in log-phase (2% glucose), AGR, and AGR+0.1% oleate (OA), and AGR+15min 40°C. **C**) Manders M1 coefficient of Erg6-mNg colocalization with LD stain MDH in various conditions. **D**) Pln1/Pet10-mNg in log and 4hrs AGR. **E**) M1 coefficient of Pln1-mNG with LD targeting. Statistics are one-way ANOVA. Scale bars 5μm.

To more directly test whether the biophysical properties of LD lipids influenced Erg6-mNg localization, rather than other metabolic changes attributed to AGR stress, we briefly heated Erg6-mNg expressing yeast after 4hrs AGR to 40°C, which is above the predicted phase transition temperature for smectic-phase SEs. Indeed, Erg6-mNg significantly, although not fully, re-localized from the ER network to LDs after only 15 minutes at 40°C (**Figure 2B**). To quantify the extent of Erg6-mNg LD localization, we calculated its relative Manders M1 coefficient, which measures total Erg6-mNg signal that overlaps with LD marker MDH. 4hrs AGR stress was accompanied by an ~75% decrease in Erg6-mNg positive LDs (**Figure 2C**). In agreement with imaging, addition of 0.1% OA returned the M1 coefficient to WT values. Brief heating also significantly, though not fully, increased the M1 coefficient.

Next, we investigated whether AGR caused a general de-localization of other canonical LD proteins from LDs. However, Pln1-mNg, a perilipin-like protein also known as Pet10 (Gao et al., 2017), maintained stable LD association following 4hrs AGR, suggesting the de-localization of LD proteins during LCL-LD formation may be selective (**Figure 2D, E**). Recently, perilipin homo-oligomerization was proposed to contribute to the stable association of perilipins on LDs (Giménez-Andrés et al., 2021). To test whether oligomerization could enhance LD protein targeting during AGR, we artificially oligomerized Erg6 by tagging it with tetrameric DsRed2. Indeed, unlike monomeric Erg6-mNg, Erg6-DsRed2 maintained LD targeting during 4hrs AGR (**SFigure 2B**). Collectively, this suggests that: 1) LD protein de-localization during AGR-associated LCL-LD formation may be selective for certain proteins, and 2) oligomerization may enhance protein retention on these LDs.

### Imaging known LD proteins reveals their selective retargeting to the ER during AGR

Given the different targeting patterns of Erg6 and Pln1 in AGR, we next examined the location of other annotated LD proteins by tagging them with mNeonGreen (mNg) and examining them in log-phase and 4hrs AGR-treated yeast. As expected, four known LD proteins Rer2-mNg, Hfd1-mNg, Yeh1-mNg (an LD-localized SE lipase), mNg-Say1 (which is annotated to target both LDs and the ER network), primarily decorated LDs in log-phase yeast. However, after 4hrs AGR all four proteins displayed ER and nuclear envelope localization, and displayed significantly reduced M1 coefficients, like Erg6 (**Figure 3A, B**). Similarly, Ayr1-mNg (a bifunctional lipase), as well as Anr2-mNg (a LD protein of unknown function predicted to be palmitoylated) also localized to LDs in log-phase yeast, but displayed primarily ER network targeting after 4hrs AGR (**SFigure 3A**). Collectively, this suggests that similar to Erg6, many canonical LD proteins exhibit more ER localization following AGR exposure, and indicates that LCL-LD formation may alter the protein composition of the LD surface.

**Figure 3:**
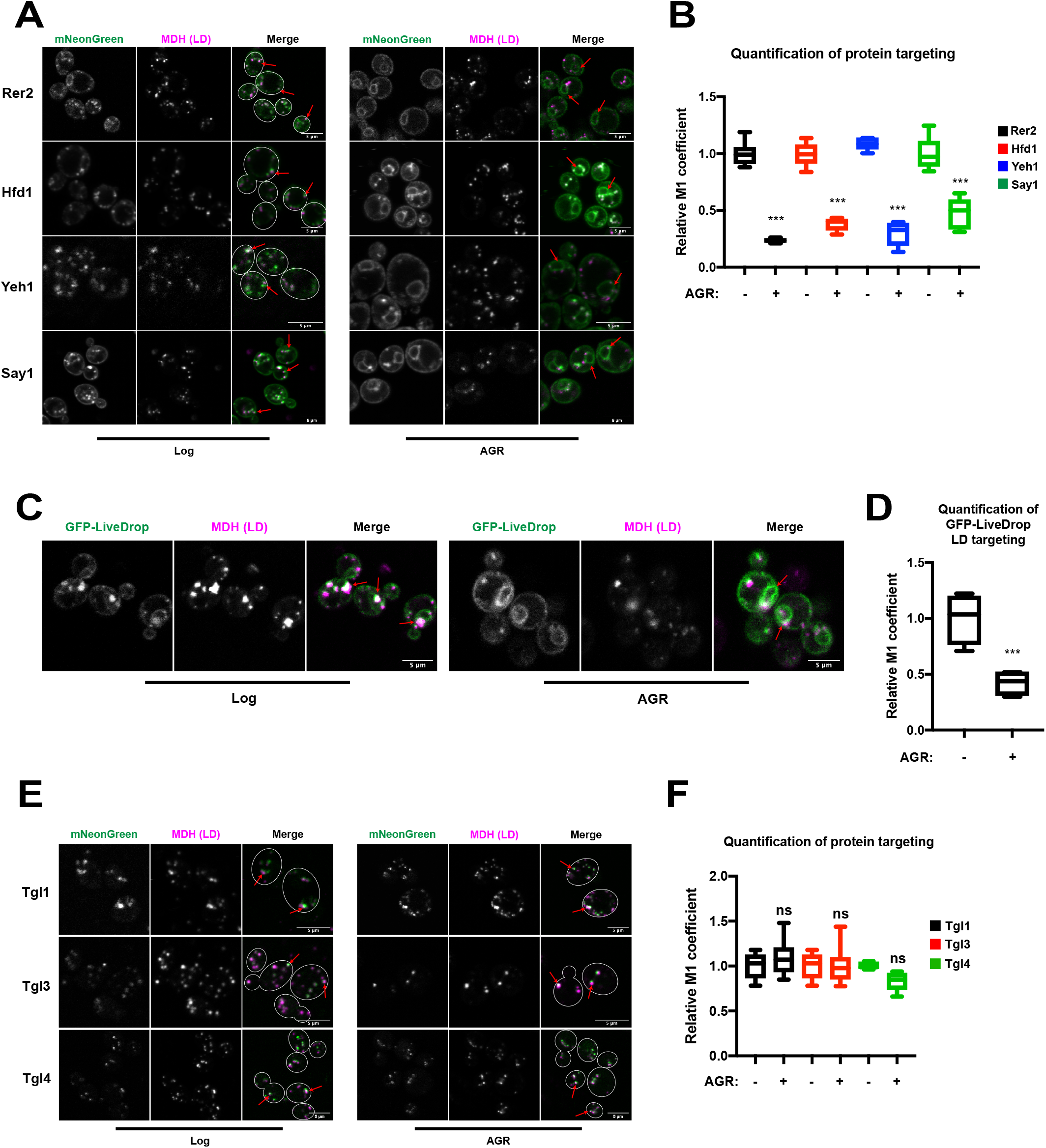
Fluorescence imaging reveals selective remodeling of LD proteome during AGR. **A**) Yeast with mNeongreen (mNg)-tagged LD proteins with MDH LD stain in log and 4hrs AGR. **B**) M1 coefficient of proteins in A. **C**) Yeast with GFP-LiveDrop and MDH LD stain in log and 4hrs AGR yeast. **D**) M1 coefficient of proteins in C. **E**) Yeast with mNg-tagged Tgl1,3,4 and stained with MDH LD marker in log-phase or 4hrs AGR. **F**) M1 coefficient of proteins in E. Scale bars 5μm.

Protein movement between the LD and ER compartments has previously been described for Type I LD proteins, which move between the ER and LDs via lipidic bridges connecting them (Wang et al., 2016). Although we observed several proteins that localized more prominently to the ER versus LDs during AGR, whether any of these represented canonical Type I LD proteins was not clear. Therefore, to interrogate whether Type I LD proteins could be re-targeted or retained at the ER during LCL-LD formation, we monitored GFP-tagged LiveDrop (Wang et al., 2016), a minimal model polypeptide for Type I LD proteins, in log-phase and 4hrs AGR-treated yeast. As expected, GFP-LiveDrop localized predominantly to LDs in log-phase yeast, but a dim ER network signal was also detected, consistent with its dual organelle targeting (**Figure 3C**). In contrast, following 4hrs AGR GFP-LiveDrop was more prominently at the ER network, and its M1 coefficient was significantly decreased (**Figure 3C, D**). This suggests that AGR and the associated LCL-LD formation promotes Type I LD protein re-distribution to, or retention at, the ER network versus LDs.

Since TG lipases were required for LCL-LD formation in AGR (**Figure 1G, M**), we next monitored the sub-cellular localization of all Tgl lipases by fluorescence microscopy. As expected, the major TG lipase Tgl3-mNg, as well as Tgl4-mNg (TG lipase) and Tgl1-mNg (SE lipase) all decorated LDs in log-phase yeast (**Figure 3E, F**). Remarkably, all three proteins retained LD localization following 4hrs AGR, likewise displaying unaltered M1 coefficients (**Figure 3E, F**). Tgl5-mNg (TG lipase) also displayed LD targeting in both log-phase and 4hrs AGR yeast (**SFigure 3B**). This suggests that in contrast to several other LD proteins, Tgl lipases maintain LD association during AGR, where they locally deplete the LD TG pool, promoting lipid phase transitions within the LD.

### Comparative proteomics reveals changes to the LD proteome in AGR stress

Since fluorescence imaging revealed that several LD proteins change sub-cellular distribution in AGR conditions, we next aimed to comprehensively map how AGR stress alters the LD proteome. We performed LC-MS/MS proteomics on LDs that were isolated from log-phase and 4hrs AGR-treated yeast using density gradient centrifugation (**Figure 4A**). To evaluate the quality of our LD isolation protocol, we performed Western blotting of whole-cell lysates and the subsequent LD isolation fractions. We found a clear de-enrichment of mitochondrial protein Por1 and the abundant plasma membrane protein Pma1 in the LD fractions, suggesting the LD fractions were relatively pure (**Figure 4B**).

**Figure 4:**
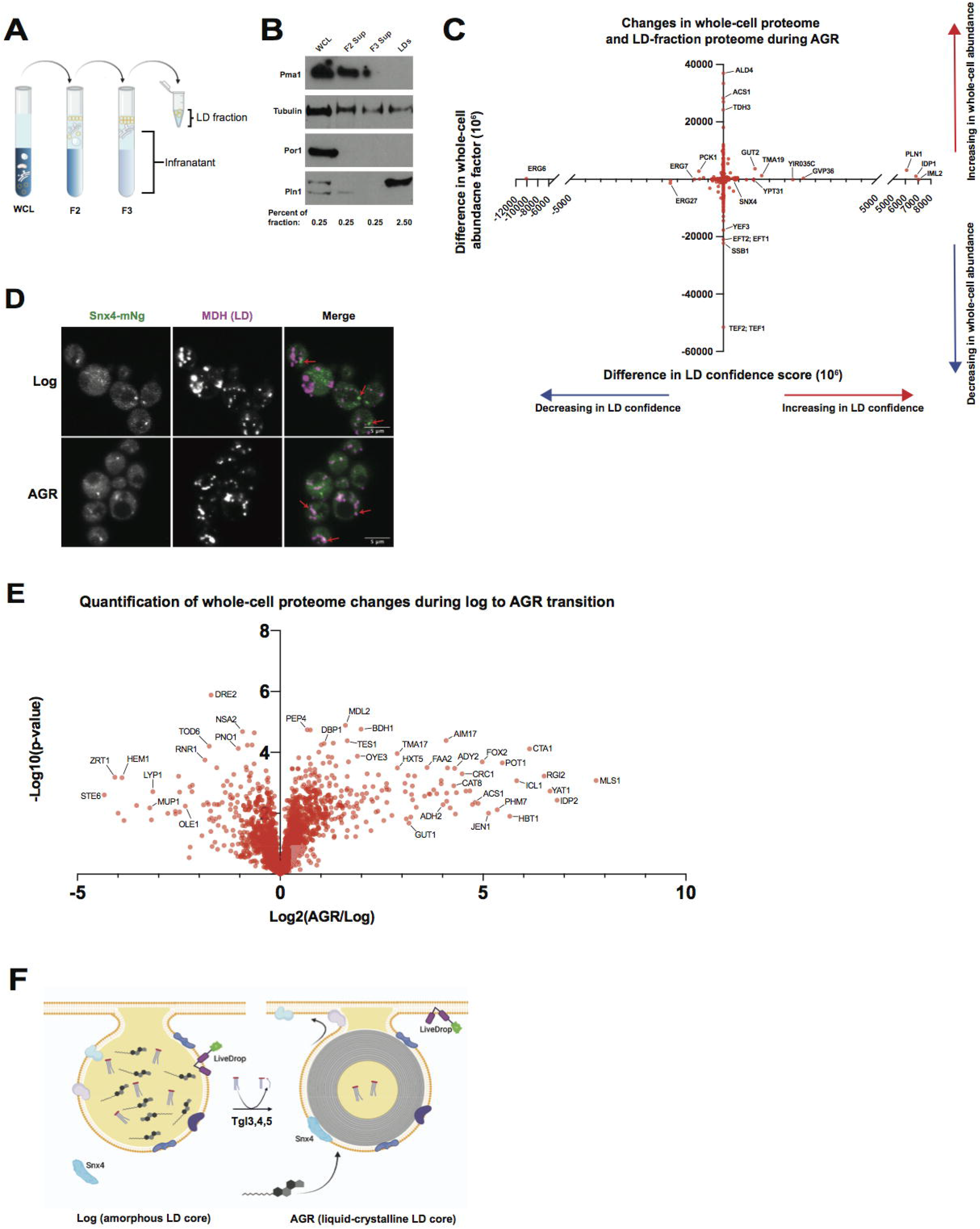
Comparative proteomics indicates non-canonical protein association with LDs, and metabolic remodeling during AGR. **A**) Schematic of LD isolation protocol. **B**) Western blot of whole cell lysate (WCL), and fractions of LD isolation protocol as in A. Pma1: plasma membrane marker, Por1: mitochondria marker. Pln1: LD marker. Tubulin: cytoplasmic marker **C**) Plot of protein abundances in whole-cell proteomics (y-axis) versus their change in LD confidence score (see methods for description of this value) Data are average of 4 independent expts. **D**) Micrographs of Snx4-mNg and LD/MDH stain in log and 4hrs AGR yeast with M1 coefficient of LD colocalization. **E**) Volcano plot showing log_10_ p-value and log_2_ abundance changes in whole-cell abundance of proteins in 4hrs AGR treatment versus 2% glucose log-phase growth. Proteins on right are increased in whole-cell abundance with 4hrs AGR, those of left decreased in abundance. Data are average of 4 independent expts. **F**) Model depicting TG lipolysis driven LCL-LD formation, and resulting changes in LD translocation to ER network targeting. Scale bars 5μm.

Given that AGR stress likely changes the global abundance of some proteins, we also conducted LC-MS/MS proteomics on the non-LD infranatant fractions generated during LD isolation, as well as whole-cell lysates of yeast in log-phase or 4hrs AGR treatment. We combined these datasets with our isolated LD proteomics to obtain a more robust dataset of high-confidence LD proteins in these conditions. This approach generated an adjusted LD enrichment score, defined as the “LD confidence score”. The approach is based on previous work from (Bersuker et al., 2018), and accounts for the spectral abundance of each protein in the LD fraction, while subtracting out the corresponding abundance from the non-LD infranatant fraction. Plotting this LD confidence score (x-axis) as a function of protein whole-cell abundances (y-axis) thus identified candidate proteins that enriched or de-enriched in AGR-associated LD fractions (**Figure 4C**). For example, proteins that increased in relative abundance in LD fractions during AGR are represented on the right side of the x-axis, whereas those that decreased are on the left side. It should be noted that many proteins did not change greatly in overall whole-cell abundance, and are thus are positioned along 0 on the y-axis. As expected, many proteins not normally associated with LDs change little on the x-axis, but may change substantially in whole-cell abundance during AGR, and are thus positioned vertically along the y-axis.

As expected, this approach revealed that Erg6 was among the most de-enriched proteins in LD fractions at 4hrs AGR (**Figure 4C**, **left side of plot**), whereas Pln1 was one of the most enriched (**Figure 4C**, **right side of plot**). Notably the LC-MS/MS detected nearly all annotated LD proteins (Currie et al., 2014), although some of these displayed changes in abundance that appeared different from the localization patterns we observed by fluorescence microscopy (**SFigure 4A**). The reason for these distinctions likely reflects the differences between imaging and biochemical methodologies, as well as some (expected) contamination of the LD fractions with co-purifying ER membranes during the LD isolation.

Using this approach, our proteomics also revealed a subset of proteins that are not annotated to localize to LDs, but were nonetheless detected in high abundance in the isolated LD fractions during AGR stress. This included Iml2 (**Figure 4C, right side of plot**), which is a sterol-associated protein required for the clearance of protein inclusions, and was previously observed associated with LDs bound to inclusion bodies (Moldavski et al., 2015). To investigate this, we imaged mNg-tagged Iml2, revealing that Iml2-mNg was throughout the cytoplasm in log-phase yeast, whereas it subtly decorated the nuclear envelope and cortical ER at 4hrs AGR (**SFigure 4B**). Even though we did not visibly detect Iml2-mNg on LDs, this may be because LDs need to be associated with protein inclusions for Iml2 to visibly enrich on them by fluorescence microscopy (Moldavski et al., 2015).

Our proteomics also indicated that two proteins containing Bin/Amphiphysin/Rvs (BAR) domains involved in Golgi/endosomal membrane trafficking, Snx4 and Gvp36, were enriched on LDs following 4hrs AGR (**Figure 4C**). BAR domains are membrane binding modules, and many BAR proteins contain amphipathic helices or other membrane inserting modules that could, in principle, insert into LDs. Furthermore, BAR protein GRAF1a was previously observed on LDs in human cells (Lucken-Ardjomande Häsler et al., 2014). Indeed, while Snx4-mNg formed cytoplasmic foci not colocalized with LDs in log-phase growth, Snx4-mNg foci did appear co-localized with a subset of LDs following 4hrs AGR (**Figure 4D**). In contrast, Gvp36-mNg distributed mostly throughout the cytoplasm in both log-phase and 4hrs AGR stress, and was not detectably enriched on LDs by fluorescence microscopy (**SFigure 4B**). Collectively, this indicates that AGR stress, which results in SE phase transition and LCL-LD formation, also selectively remodels the LD proteome. The uncoating of canonical proteins from LDs may lead to enhanced LD association of non-canonical factors or membrane trafficking proteins with the phospholipid surface of LDs.

### Global proteomics indicates AGR promotes fatty acid oxidation during metabolic remodeling

Energy depletion drives metabolic remodeling in yeast, favoring the reorganization of organelles and the utilization of alternative carbon sources when glucose is restricted (Marini et al., 2020) (Eisenberg and Büttner, 2014). Since we conducted whole-cell LC-MS/MS proteomics of log-phase and 4hrs AGR yeast, we next examined these datasets to determine whether changes in whole-cell protein abundances revealed patterns of metabolic remodeling that involved LDs and their lipids. Indeed, we found that 4hrs AGR stress induced changes in the abundances of many proteins involved in fatty acid metabolism. In particular, peroxisome enzymes involved in fatty acid oxidation (FAO), including Pot1, Fox2, and Cta1 were among the most increased in abundance during AGR compared to log-phase growth (**Figure 4E, right side of plot**). Also elevated were the peroxisome-associated fatty acyl-CoA ligase Faa2, the acetyl-CoA transporter Crc1 (which transports acetyl-CoA derived from peroxisome FAO to mitochondria), as well as Yat1, a carnitine acetyl-transferase that works with Crc1 to promote acetyl-CoA utilization within mitochondria. Enzymes related to the tricarboxylic acid cycle including Icl1 and Idp2, the malate synthase Mls1, and acetyl-CoA synthase Acs1 were also among the most elevated proteins in AGR-treated yeast (**Figure 4E**). In contrast, amino acid transporters like Mup1 and Lyp1 were significantly decreased in abundance (**Figure 4E, left side of plot**), consistent with their turnover during glucose starvation that promotes adaptive metabolic remodeling (Lang et al., 2014), (Wood et al., 2020).

Collectively, this indicates that glucose restriction promotes the mobilization of TGs from LDs that may provide fatty acids as fuel for cellular energetics in peroxisomes and mitochondria. Indeed, acetyl-CoA generated by peroxisome FAO can be delivered to mitochondria to fuel its energetics in the absence of glucose, suggesting inter-organelle remodeling during glucose restriction that enables LD-derived lipids to ultimately fuel alternative carbon metabolism. An additional consequence of this Tgl-dependent TG mobilization is a shift in the neutral lipid ratios in LDs, ultimately giving rise to SE transition into a liquid-crystalline phase within the LDs.

## Discussion

Emerging evidence suggests the phase transition properties of cellular biomolecules, such as proteins in membraneless organelles, directly influence cell physiology and organization. Like proteins, lipids also undergo phase transitions, and can form liquid-crystalline lattices that are observed in human diseases like atherosclerosis, or in organelles like LDs. However, the metabolic cues that drive these phenomena, and their impact on organelle physiology, are unclear. Here we show that in yeast, AGR stress promotes the formation of liquid-crystalline lipid phase transitions within LDs. These transitions require TG lipolysis, suggesting the loss of TG within the hydrophobic core of LDs promotes the transition of SEs from an amorphous to a smectic liquid-crystalline phase. In agreement with this, we find AGR drives metabolic remodelling that elevates peroxisome-mediated lipid oxidation. Furthermore, we provide evidence that LCL-LD phase transitions alter the LD proteome (**Figure 4F**).

How proteins are targeted to LDs is still poorly understood, and involves trafficking from the ER network or cytoplasm to the LD surface. In this study, we revealed that the LD proteome dramatically differs between AGR-treatment and log-phase growth. Erg6, a canonical LD protein, relocalizes to or is retained at the ER network, suggesting it moves from LDs to the ER via a lipidic bridge. This LD delocalization appears suppressed or quickly reversed when yeast cells are briefly heated to 40°C, (i.e. above the predicted melting temperature of smectic-phase SEs), suggesting direct movement of the proteins between LD and ER via ER-LD connections. In line with this, GFP-LiveDrop, which under log-phase conditions targets primarily to LDs, appears predominantly ER localized during AGR. Collectively, this suggests that Type I LD proteins favor ER localization versus the surface of LCL-LDs. This also indicates that many yeast LDs maintain connections to the ER network and thus exhibit the lipidic bridges necessary for this inter-organelle trafficking, consistent with earlier work (Jacquier et al., 2011). The redistribution of LD proteins to the ER may be due to changes in LD monolayer fluidity after LCL-LD formation, which could alter the energetic favorability of proteins to remain on the LD surface. We also cannot rule out that the lipid composition of the ER network changes during AGR to a state that favors protein targeting or retention. We also find that artificially multimerizing Erg6 with a DsRed2 tag promotes its LD retention at AGR, implying protein oligomerization enhances LD retention, as has previously been observed for perilipins (Giménez-Andrés et al., 2021).

Whereas Erg6 delocalized from LDs during AGR, TG lipases Tgl3,4,5 remained LD bound. Although the LD anchoring mechanisms for Tgl lipases are not fully understood, this implies that LDs continue to mobilize TG during AGR, gradually altering the TG:SE neutral lipid ratio in a manner that supports SE phase transition. Indeed, AGR-treated yeast contain less TGs, consistent with lipolysis that provides fatty acids to fuel metabolic energetics. Fatty acids derived from these TGs are likely substrates for peroxisome FAO, of which several key enzymes are elevated during AGR stress. The acetyl-CoA produced from FAO could also fuel mitochondrial energetic pathways, several proteins of which are elevated by proteomics. LCL-LDs also exhibited de-targeting of enzymes like Hfd1, Rer2, and Say1. It is possible these enzymes’ re-distributions influences their activities, and therefore promote metabolic remodeling. Indeed, several Erg pathway enzymes also appeared de-enriched from LDs during AGR by proteomics, and Erg1 is more active at the ER than on LDs (Leber et al., 1998).

Our proteomic and imaging analysis also revealed that LDs may become decorated with non-LD proteins during AGR stress. This included the BAR domain protein Snx4, which co-localized with some LDs only during AGR stress. As BAR proteins contain membrane binding/inserting modules, it is possible that Snx4 associates with LDs during AGR by inserting into its monolayer surface. Since the LD surface is normally densely coated with proteins, it is also possible Snx4 and other proteins may associate with the LD surface as it is uncoated of canonical LD proteins during AGR stress. Proteomics also detected Iml2 on LDs during AGR. Previous work proposed that Iml2 associated with LDs, and promoted the delivery of sterols to protein inclusions during their clearance in an unknown mechanism involving LDs (Moldavski et al., 2015). Although unclear, it is possible Iml2 may influence sterol metabolism on LCL-LDs.

This study is a significant step toward enhancing understanding how lipid phase transitions influence LD and organelle protein composition and ultimately function. Future studies will interrogate whether such changes in the LD proteome reflect metabolic remodeling that ultimately enable yeast to adapt to glucose shortage.

## Materials and Methods

Please see STAR Methods for a full description of the Methodology.

## Supporting information

SFigure 1

SFigure 2

SFigure 3

SFigure 4

SMovie 1

SMovie 2

SMovie 3

## Acknowledgement

We thank Jonathan Friedman, and members of the Henne and Nicastro labs for helpful insights during this study. We thank Daniel Stoddard for management of the UTSW electron microscope facilities and training, and Gang Fu for some data acquisition. The UT Southwestern Cryo-Electron Microscopy Facility is supported in part by the CPRIT Core Facility Support Award RP170644. We would also like to thank the UTSW proteomics and live cell imaging facilities for their assistance with data collection and analysis. Finally, we would like to thank Dr. Joel Goodman for the Pln1 antibody. W.M.H. is supported by funds from the Welch Foundation (I-1873), the NIH NIGMS (GM119768), NIDDK (DK126887), Ara Parseghian Medical Research Fund, and the UT Southwestern Endowed Scholars Program. S.R. is supported in part by a NIH T32 training grant (5T32GM008297). L.G., E.R., and D.N. are supported by the Cancer Prevention and Research Institute of Texas grant RR140082 to D.N. This research was supported in part by the computational resources provided by the BioHPC supercomputing facility located in the Lyda Hill Department of Bioinformatics, UT Southwestern Medical Center.

## Supplemental Figure Legends

**Supplemental Figure 1: LD lipid phase transitions characterized by cryo-FIB and cryo-ET. A-J**) Electron dose series (“bubblegrams”) for LDs from cryo-FIB milled WT yeast in log phase (**A-E**) or after AGR (**F-J**); series of 2D cryo-EM images were recorded of the same LDs exposed to increasing electron dose (1 - 400 e-/Å2). Note that liquid-crystalline layers (LCLs) (see box in **G** magnified in **J**) and excessive bubbling in LD centers (starting at an electron dose <30 e-/Å^2^) occurred only under AGR. Even at 400 e-/Å^2^ electron dose, minimal bubbling (white arrowheads in **E**).was observed in log WT. **K-M**) Representative tomographic slices from cryo-FIB-milled and cryo-ET reconstructed WT in log phase (**K**), *nvj1△* yeasts after 4hrs AGR (**L**), and *are1are2△* yeast after 4hrs AGR (**M**). The nucleus-vacuole junction (black arrowheads in **K** and **M**) was observed in WT and *are1are2Δ* yeasts, but absent in *nvj1△* yeast (white arrowhead in **L**). No LDs were found in *are1are2△* yeast. V, vacuole. N, nucleus. L, lipid droplet, M, mitochondrion. **N**) Yeast stained with LD marker MDH in log and 4hrs AGR. Scale bars: 50nm (A-I), 200nm (K-M), 25nm (J).

**Supplemental Figure 2: Erg6 LD targeting is influenced by AGR-associated LCL-LD formation**. **A**) WT or *tgl3,4,5Δ* Erg6-GFP yeast in log or AGR conditions. **B**) Erg6-DsRed2 localized to LDs in log and 4hrs AGR. Scale bars 5μm.

**Supplemental Figure 3: Selective delocalization of LD proteins during AGR stress. A**) Yeast expressing mNg-tagged Ayr1 and Anr2 and stained with LD marker MDH in log-phase or 4hrs AGR conditions. **B**) Yeast expressing Tgl5-mNg with MDH LD stain. Scale bars 5μm.

**Supplemental Figure 4: Additional LD proteins examined in log-phase and AGR conditions**. **A**) Heat map depicting relative % changes in annotated LD proteins from log to 4hrs AGR conditions. Average of 4 independent log-phase and 4hrs AGR experiments. **B**) Yeast with mNg-tagged Iml2 or Gvp36 and stained with MDH in log-phase and 4hrs AGR conditions. Scale bars 5μm.

**Supplemental Movie 1**: Tomographic reconstruction of a cryo-FIB-milled WT yeast cell after 4hrs acute glucose restriction. Compare with Figure 1A. Scale bar: 200nm.

**Supplemental Movie 2**: Tomographic reconstruction of a LD from a cryo-FIB-milled WT yeast cell in log phase (grown with 2% glucose). Compare with Figure 1B. Scale bar: 50nm.

**Supplemental Movie 3**: Tomographic reconstruction of a LD from a cryo-FIB-milled WT yeast cell after 4hrs AGR. Compare with Figure 1C. Scale bar: 50nm.

