## Supplementary figures and images for "Liquid-crystalline lipid phase transitions in lipid droplets selectively remodel the LD proteome"

### SFigure 1

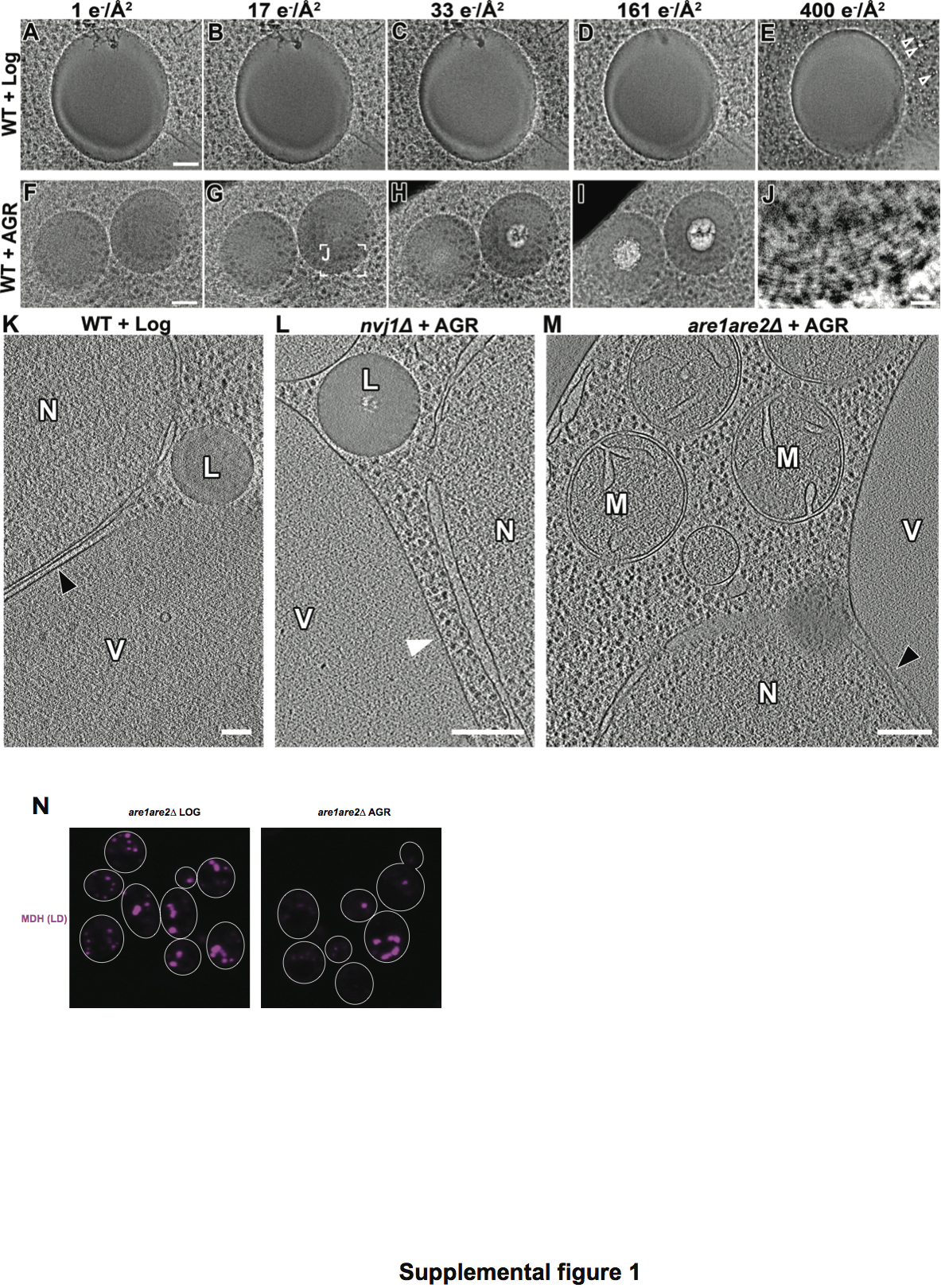

### SFigure 2

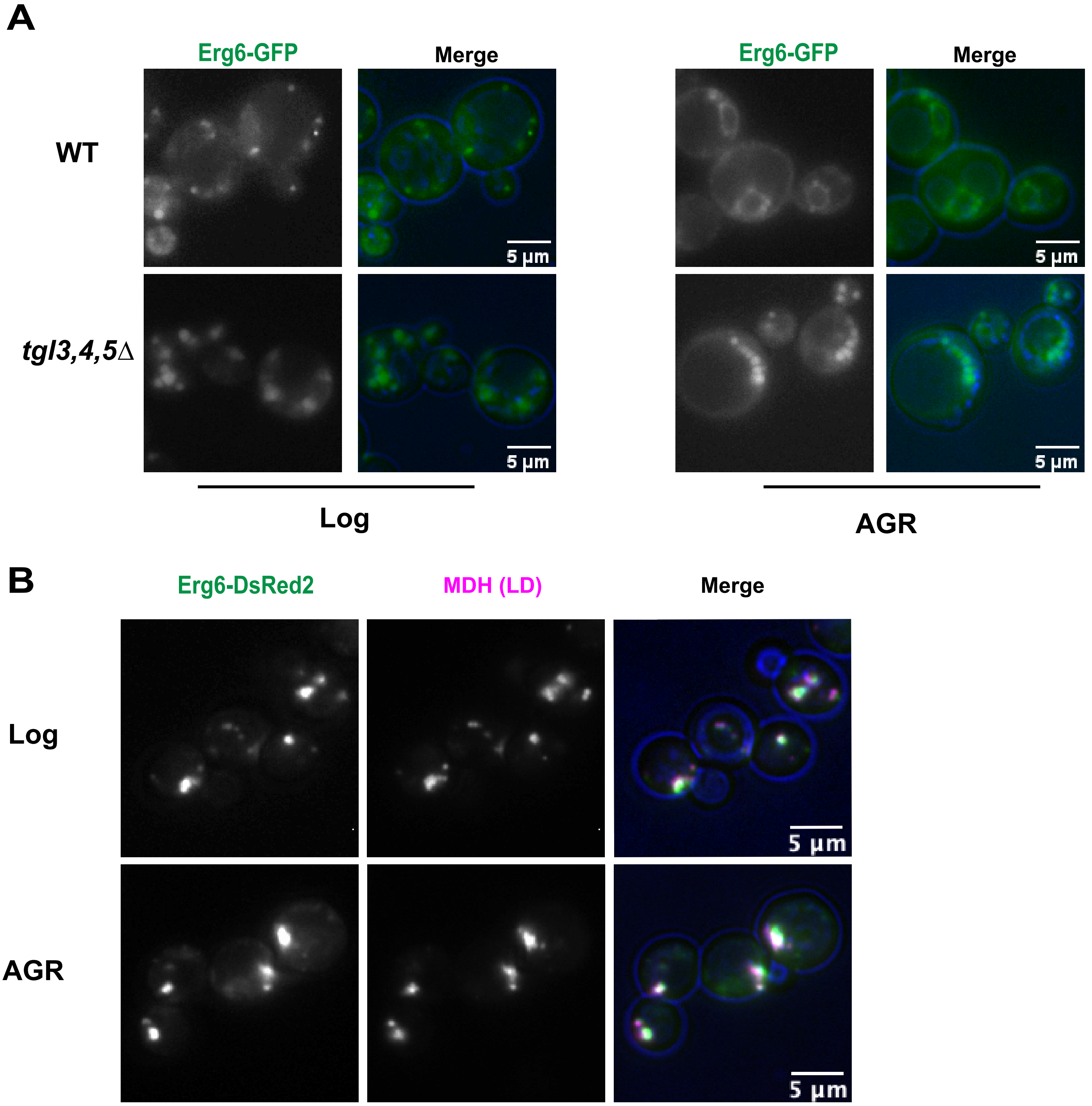


Supplemental figure 2

### SFigure 3

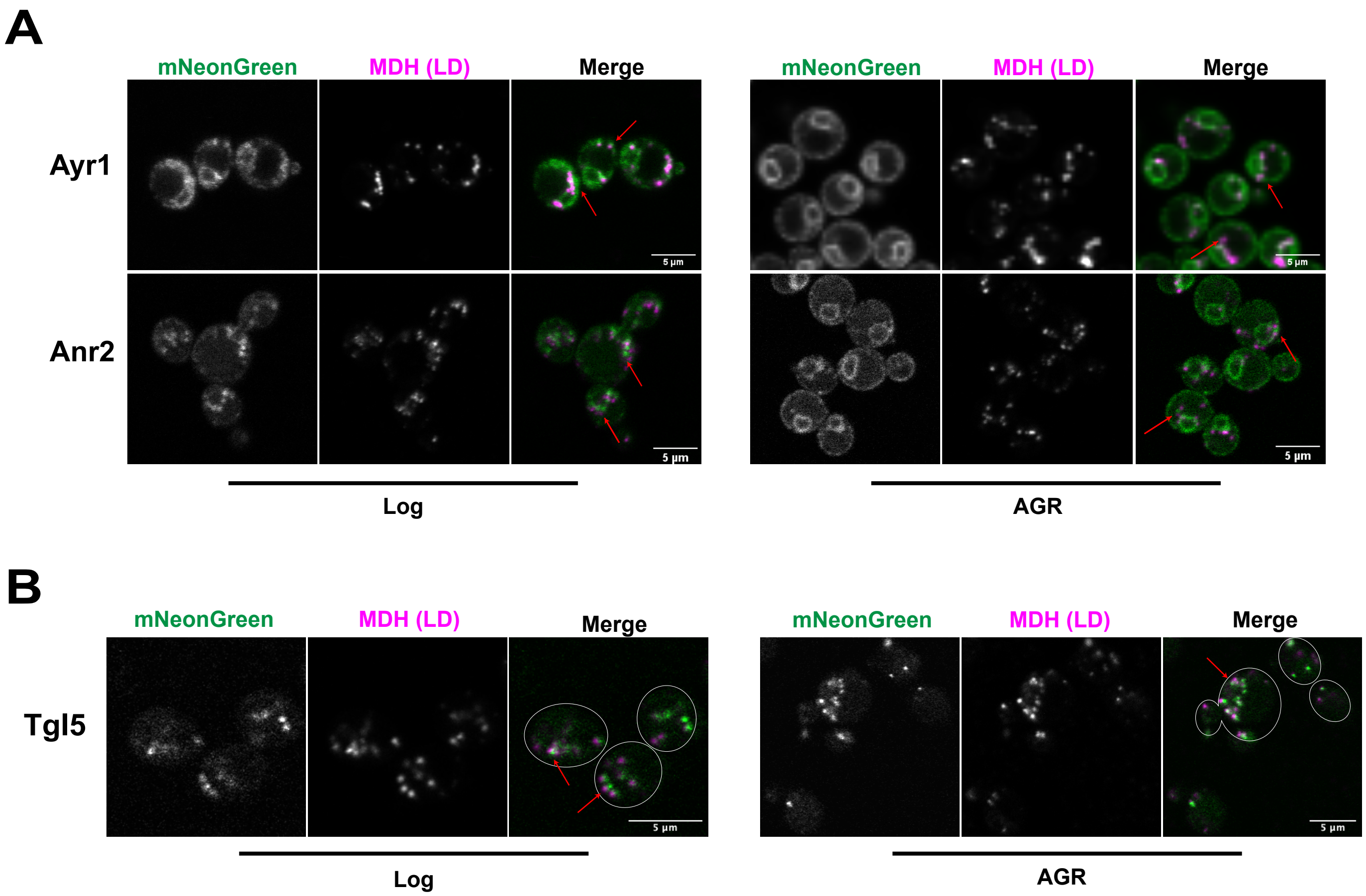


Supplemental figure 3

### SFigure 4

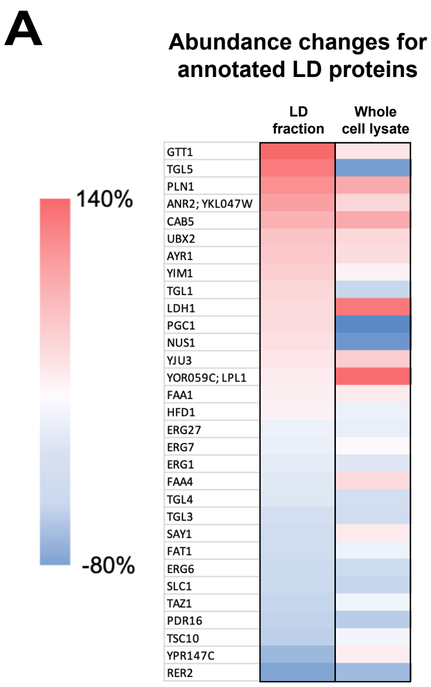


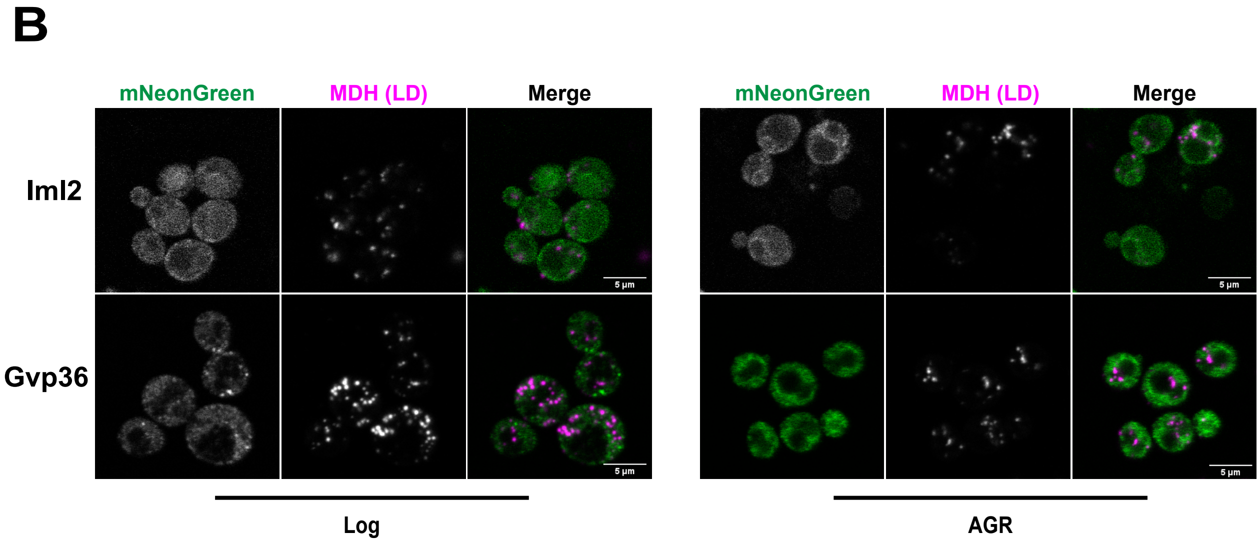


**Supplemental figure 4**
