## Supplemental Methods for "Liquid-crystalline lipid phase transitions in lipid droplets selectively remodel the LD proteome"

**Star Methods for Rogers, et al.**

**KEY RESOURCES TABLE**

| REAGENT or RESOURCE | SOURCE | IDENTIFIER |
| --- | --- | --- |
| Chemical, peptides, and Recombinant Proteins |  |  |
| Nourseothricin sulfate (NAT) | Gold Biotechnology | Cat# N5001 |
| Hygromycin B | Sigma-Aldrich | Cat# 10843555001 |
| Yeast Nitrogen Base Without Amino Acids | Sigma-Aldrich | Cat# Y0626 |
| Yeast synthetic drop out medium supplements – without uracil | Sigma-Aldrich | Cat#Y1501 |
| Zymolyase 20T | Amsbio | Cat# 120491-1 |
| Phusion DNA Pollymerase | NEB | Cat# M0530 |
| Mg132 in DMSO | Sigma | Cat# M7449 |
| 4x NuPAGE LDS sample buffer | ThermoFisher | Cat# NP0008 |
| Monodansylpentane (MDH) - Autodot | Abcepta | Cat# SM-1000a |
| Critical Commercial Assays |  |  |
| Gibson assembly mastermix | NEB | Cat# E2611 |
| ORGANISMS/STRAINS | **SOURCE** | **IDENTIFIER** |
| Saccharomyces cerevisiae W303 (leu2-3,112 trp1-1 can1-100 ura3-1 ade2-1 his3-11,15) | Saccharomyces GENOME DATABASE https://www. yeastgenome.org/strain/W303 | N/A |
| W303 Erg6-mNg | This paper | N/A |
| W303 *tgl3,4,5*Δ | This paper | N/A |
| W303 *nvj1*Δ | Rogers, ELife, 2021 | N/A |
| W303 *are1are2*Δ | Gao et al. JCB, 2017 | N/A |
| W303 Pln1-mNg | This paper | N/A |
| W303 Erg6-DsRed2 | This paper | N/A |
| W303 Rer2-mNg | This paper | N/A |
| W303 Hfd1-mNg | This paper | N/A |
| W303 Yeh1-mNg | This paper | N/A |
| W303 pRS305::ADH::mNg-Say1::LEU | This paper | N/A |
| W303 pBP73G::GFP-LiveDrop::URA | This paper | N/A |
| W303 Ayr1-mNg | This paper | N/A |
| W303 Tgl5-mNg | This paper | N/A |
| W303 Anr2-mNg | This paper | N/A |
| W303 pRS305::ADH::mNg-Gtt1::LEU | This paper | N/A |
| W303 Tgl1-mNg | This paper | N/A |
| W303 Tgl3-mNg | This paper | N/A |
| W303 Tgl4-mNg | This paper | N/A |
| W303 Snx4-mNg | This paper | N/A |
| W303 Iml2-mNg | This paper | N/A |
| W303 Gvp36-mNg | This paper | N/A |
| RECOMBINANT DNA | **SOURCE** | **IDENTIFIER** |
| pBP73G (*URA3* selection plasmid with GPD promoter on pRS416 background) | Scott Emr lab | N/A |
| pRS305 (*LEU2* selection plasmid with ADH promoter) | Scott Emr lab | N/A |
| Software and Algorithms |  |  |
| Fiji (ImageJ) for Relative M coefficient | https://imagej.nih.gov/ij/ | N/A |

**LEAD CONTACT AND MATERIALS AVAILABILITY**

Further information and requests for resources and reagents should be directed to and will be fulfilled by the Lead Contact, W. Mike Henne. Requests will be handled according to the University of Texas-Southwestern policies regarding MTA and related matters.

**METHOD DETAILS**

**Yeast growth conditions**

The wild-type parental strain used for all experiments and cloning in this study was W303 (leu2-3, 112 trp1-1 can1-100 ura3-1 ade2-1 his3-11,14). Synthetic-complete (SC) growth media was used for culturing yeast cells in all experiments, except for experiments where uracil was excluded to accommodate the retention of pBP73-G plasmids. For all experiments, a streak of yeast was inoculated from a YPD (yeast extract peptone dextrose) plate into SCD (synthetic complete dextrose/glucose) media and allowed to grow overnight in a 30°C incubator with shaking at 210RPM. Overnight cultures were diluted to an OD_600_=0.2 in SCD media containing 2% glucose (w/v). Log-phase yeast were collected at OD_600_=0.5. For AGR (acute glucose restriction) treatment, OD_600_=0.5 yeast were pelleted by centrifugation, briefly washed with SC media containing no glucose, and resuspended in SCD media containing specifically dextrose/glucose at a final concentration of 0.001% (w/v). AGR-treated cells were grown at 30°C with 210RPM shaking for the indicated times. Where indicated, cells were also treated with 0.1% oleic acid (Sigma) or 50μM Mg132 (M8699, Sigma; proteasome inhibitor).

**Strain generation and plasmid construction**

The established lithium acetate method was used for the generation of all yeast knockouts and knockins. Briefly, yeast were diluted from an overnight culture to an OD_600_=0.2 in YPD media until they reached OD_600_=0.6. For each transformation, approximately 7.5OD units of cells were pelleted, washed with 0.1M lithium acetate, pelleted again, and resuspended in 700μL of transformation solution (40% PEG in 0.1M lithium acetate, 0.25μg/μL single stranded carrier DNA (D9156, Sigma)) supplemented with 50μL of PCR product. Transformations were kept at room temperature for one hour in transformation solution. Immediately prior to heat shock, DMSO was added to a final concentration of 10% (v/v). Cells were heat shocked at 42°C for 15 minutes, followed by 1 minute incubation on ice and subsequent plating onto YPD solid media. Cells were recovered overnight on YPD plates in a 30°C incubator and replica plated onto either YPD or SC selective solid media the following day. Plasmids were generated for this study using Gibson Assembly following the manufacturer’s protocol (E2611, NEB). All pBP73-G vectors were cut with XbaI and XhoI.

**Cryo-sample preparation and cryo-FIB milling**

4 μl of the cultured yeast cells were added to a glow-discharged (30 seconds at -30 mA) copper R2/2 holey carbon grid (Quantifoil Micro Tools GmbH, Jena, Germany), then the grid was rapidly plunge-frozen in liquid ethane using a homemade plunge freezer device and stored in liquid nitrogen until used. Cryo-FIB milling was performed as previously described (Hariri et al., 2019). Briefly, grids were mounted in notched cryo-FIB Autogrids (Thermo Fisher Scientific, MA, USA), then loaded into a shuttle under cryogenic conditions and transferred into an Aquilos dual-beam instrument equipped with a cryo-stage (FIB/SEM; Thermo Fisher Scientific). The sample surface was sputter-coated with platinum for 20 s at 30 mA current and then coated with a layer of organometallic platinum using the gas injection system for 6s at a distance of 1 mm before milling. The stage was then tilted to 10°-18° (so that the bulk-mill-holes lined up in front and behind the cell), and the cell was milled with 30 kV gallium ion beams of 100 pA current for rough milling and 10 pA for polishing until the final lamella was 100-200nm thick.

**Cryo-ET and image processing**

Lamellae were imaged using a Titan Krios transmission electron microscope (FEI/Thermo Fisher Scientific) operated at 300kV. Images were captured using either a 4k × 4k K2 direct electron detection camera (Gatan, Pleasanton, CA) at a magnification of 26,000x (5.5 Å pixel size) or a 5k × 6k K3 direct electron detection camera (Gatan, Pleasanton, CA) at a magnification of 15,000x (5.7 Å pixel size), which were located behind a Bioquantum post-column energy filter (Gatan) that was operated in zero-loss mode (20-eV slit width). The defocus was set to −0.5 μm using a Volta phase plate (Danev et al., 2014). Data acquisition was performed using the microscope control software SerialEM (Mastronarde, 2005) in low-dose mode. We initially recorded a montaged overview of each cryo-FIB lamella to select cellular areas of interest. Then tilt series were collected over a range of +/-56° with 2° increments using a dose-symmetric tilting scheme (Hagen et al., 2017), with the total electron dose per tilt series limited to ∼100 e/Å^2^. Counting modes of the K2 and K3 cameras were used and for each tilt image 15 frames (0.4s exposure time per frame for K2 and 0.04s exposure time per frame for K3) were recorded. The frames of each tilt series image were motion-corrected using MorionCor2 (Zheng et al., 2017) and then merged using the script extracted from the IMOD software package (Kremer et al., 1996) to generate the final tilt serial data set. Tilt series images were aligned fiducial-less using patch tracking (800 × 800-pixel size) and the tomogram was reconstructed by the back-projection method using the IMOD software package (36). For better visualization, the cryo-tomograms were denoised using nonlinear anisotropic diffusion in the IMOD package. In total, the following number of cryo-FIB lamellae per strain/condition were examined: 20 (WT + log), 17 (WT + AGR), 6 (WT + AGR + OA), 10 (*tgl3,4,5*Δ + AGR), 7 (WT + log + OA), 11 (*nvj1*Δ + AGR), and 15 (*are1are2*Δ + AGR).

**Fluorescence microscopy**

For confocal microscopy, cells were grown as described above and collected by centrifugation at 4,000xg for two minutes. Where indicated, cells were incubated for five minutes with monodansylpentane (MDH, SM1000a, abcepta) at a final concentration of 0.1mM to visualize LDs. Prior to imaging, cells were washed with 1mL of SC media and resuspended in glucose-free SC media at approximately one one-hundredth of the original volume. All images were taken as single slices at approximately mid-plane using a Zeiss LSM880 inverted laser scanning confocal microscope equipped with Zen software. Images were taken with a 63x oil objective NA=1.4 or 40x oil objective NA=1.4 at room temperature. For epifluorescence microscopy, cells were grown, stained, and collected as described above. Imaging was performed on an EVOS FL Cell Imaging System at room temperature.

**Lipid extraction and thin layer chromatography**

For lipid extraction, approximately 50OD units of cells were collected for each sample, and the pellet wet weight was normalized and noted prior to extraction. Lipid extraction was performed using a modified Folch method (Folch et al., 1957). Briefly, cell pellets were resuspended in MilliQ water with 0.5mm glass beads and lysed by three one-minute cycles on a bead beater. Chloroform and methanol were added to the lysate to achieve a 2:1:1 chloroform:methanol:water ratio. Samples were vortexed, centrifuged to separate the organic solvent and aqueous phases, and the organic solvent phase was collected. Extraction was repeated a total of three times. Prior to thin layer chromatography, lipid samples were dried under a stream of argon gas and resuspended in 1:1 chloroform:methanol to a final concentration corresponding to 4μL of solvent per 1mg cell pellet wet weight. Isolated lipids were spotted onto heated glass-backed silica gel 60 plates (1057210001, Millipore Sigma), and neutral lipids were separated in a mobile phase of 80:20:1 hexane:diethyl ether:glacial acetic acid. TLC bands were visualized by spraying dried plates with cupric acetate in 8% phosphoric acid and baking at 140°C for an hour.

**Lipid droplet isolation by density centrifugation**

The procedure for lipid droplet isolation was adapted from previously published methods in (Au - Mannik et al., 2014). Specifically, approximately 600OD units of cells were grown to appropriate growth phase in SCD media. Cells were collected by centrifugation at 4,100xg for 10 minutes at room temperature and resuspended in reducing buffer (10mM Tris-HCl pH 9.4, 10mM DTT) to a final concentration of 100OD/mL. After a five minute incubation at room temperature, cells were pelleted again by centrifugation at 4,100xg for 5 minutes. Reducing buffer was removed and replaced with equal volume spheroplasting buffer (10mM Tris-HCl pH 7.4, 700mM sorbitol, 7.5g/L yeast extract, 15g/L peptone, 1mM DTT). Spheroplasting was initiated by addition of zymolyase 20T (120491-1, AMSBIO) to a final concentration of 1mg/100OD cell units. For log-phase cells, spheroplasting buffer was supplemented with glucose to a final concentration of 0.5% (w/v), while AGR-treated cells were spheroplasted in the absence of glucose. Cells were incubated in spheroplasting buffer for 40 minutes in a 30°C incubator shaking at 130RPM. Once spheroplasting was complete, cells were pelleted by centrifugation at 4,100xg for 5 minutes at 4°C. Spheroplasting buffer was thoroughly removed and cells were gently resuspended in cold lysis buffer using a cut pipette tip (10mM Tris-HCl pH 7.4, 12% ficoll, 200μM EDTA, 1x protease/phosphatase inhibitor cocktail (78444, Thermo Fisher Scientific), 50μM Mg132, 1mM DTT) to a final concentration of 500OD/mL. Spheroplasts were once again pelleted by centrifugation at 4,100xg for 5 minutes at 4°C and resuspended in cold lysis buffer to a final concentration of 1000OD/mL and stored at -80°C.

For cell lysis, spheroplasts were thawed on ice and diluted to a concentration of 500OD/mL using cold lysis buffer. Spheroplasts were transferred to a chilled glass dounce homogenizer and lysed by 25 strokes of a loose-fitting pestle. The resulting lysate was loaded into a 11x60mm ultracentrifuge tube (344062, Beckman), overlaid with equivalent volume of lysis buffer, and centrifuged in a SW60Ti rotor at 100,000xg for 1.5 hours at 4°C set to accel=max and decel=none. Float fractions were collected using a cut P200 pipette tip and loaded into a new 11x60 ultracentrifuge tube. The bottom fraction volume was adjusted to approximately 1.5mL using cold lysis buffer and was overlaid with overlay buffer #1 (10mM Tris-HCl pH 7.4, 8% ficoll, 200μM EDTA, 1x protease/phosphatase inhibitor cocktail, 50μM Mg132, 1mM DTT). Centrifugation was repeated for 1 hour under the same settings listed above. The second float fraction was collected and transferred to a new 11x60 ultracentrifuge tube. The volume was adjusted to 1.5mL with overlay buffer #1 and supplemented with sorbitol to a final concentration of 600mM. Fractions were then overlaid with overlay buffer #2 (10mM Tris-HCl pH 7.4, 250mM sorbitol, 200μM EDTA, 1x protease/phosphatase inhibitor cocktail, 50μM Mg132, 1mM DTT) and centrifugation step was repeated for one hour. The final float fraction was collected, transferred to a 1.5mL microcentrifuge tube, and concentrated by removing the bottom infranatants following repetitive centrifugations at 20,000xg in a 4°C microcentrifuge. LD fractions were concentrated to approximately 200μL and stored at -80°C along with corresponding infranatant fractions. Note the infranatant fraction is defined here as the non-LD float fraction containing other organelles.

**Protein extractions**

***Lipid droplet fractions de-lipidation and protein extraction***

LD fractions were mixed with 1mL of -20°C 100% acetone and stored at -80°C overnight to precipitate proteins. After overnight incubation, tubes were centrifuged at 20,000xg for 10 minutes at 4°C to pellet the proteins. The supernatant was removed and the protein pellet was subjected to three washes, each followed by the same centrifugation step listed above. The washes were performed with -20°C 100% acetone, 4°C 1:1 acetone:diethyl ether, and room temperature 100% diethyl ether. After the final wash, the supernatant was removed and the protein pellet was dried in a room temperature speed vac for 15 minutes to remove residual acetone. The resultant dried pellet was resuspended in 100μL of resuspension buffer (2x NuPAGE LDS sample buffer (NP0008, ThermoFisher), 10% beta-mercaptoethanol, 8M urea). Samples were heated to 37°C for two hours accompanied by gentle mixing with a pipette prior to being subjected to SDS-PAGE.

***Whole-cell and infranatant protein extractions***

Whole cell protein extracts were isolated from 50OD units of cells. Frozen cell pellets stored at -80°C were incubated with 20% trichloroacetic acid for 30 minutes on ice with occasional mixing using a vortex. Precipitated proteins were pelleted in a 4°C centrifuge at 15,000xg for 5 minutes. After removing the supernatant, the pellet was washed three times with cold 100% acetone followed by brief sonication. After the washes, protein pellets were dried in a room temperature speed vac for 15 minutes to remove residual acetone.

For infranatant fraction analysis, 2mL of each infranatant was reserved after LD isolation and stored at -80°C. TCA was added to each infranatant fraction to a final concentration of 20%. From this point, proteins were extracted as indicated above for whole-cell lysates. Dried protein pellets from whole-cell lysates and infranatant fractions were resuspended in 200μL of resuspension buffer and heated as indicated above for LD fractions.

**Sample preparation and LC-MS/MS proteomics**

Immediately following heating, protein samples were centrifuged at 20,000xg for 5 minutes to pellet and remove insoluble materials. Each supernatant was loaded onto a 10% mini-protean TGX gel (4561033, Bio-Rad). Samples were subjected to electrophoresis at 130V constant until the dye front was approximately 10cm into the gel. The gel was subsequently removed from the casing and stained with Coomassie reagent (0.5 Coomassie G-250, 50% methanol, 10% acetic acid) for one hour on a room temperature rocker. The gel was then subjected to destaining (40% methanol 10% acetic acid) for two hours. Once the gel was sufficiently destained, 10cm gel bands were excised from each lane, taking care to exclude the stacking gel and dye front. Gel bands were further cut into 1cm squares and placed into microcentrifuge tubes. Samples were digested overnight with trypsin (Pierce) following reduction and alkylation with DTT and iodoacetamide (Sigma–Aldrich). The samples then underwent solid-phase extraction cleanup with an Oasis HLB plate (Waters) and the resulting samples were injected onto an Orbitrap Fusion Lumos mass spectrometer coupled to an Ultimate 3000 RSLC-Nano liquid chromatography system. Samples were injected onto a 75 μm i.d., 75-cm long EasySpray column (Thermo) and eluted with a gradient from 0-28% buffer B over 90 minutes. The buffer contained 2% (v/v) ACN and 0.1% formic acid in water, and buffer B contained 80% (v/v) ACN, 10% (v/v) trifluoroethanol, and 0.1% formic acid in water. The mass spectrometer operated in positive ion mode with a source voltage of 1.5-2.0kV and an ion transfer tube temperature of 275°C. MS scans were acquired at 120,000 resolution in the Orbitrap and up to 10 MS/MS spectra were obtained in the ion trap for each full spectrum acquired using higher-energy collisional dissociation (HCD) for ions with charges 2-7. Dynamic exclusion was set for 25s after an ion was selected for fragmentation. Raw MS data files were analyzed using Proteome Discoverer v 2.4 (Thermo), with peptide identification performed using Sequest HT searching against the *S. cerevisiae* protein database from UniProt. Fragment and precursor tolerances of 10ppm and 0.6Da were specified, and three missed cleavages were allowed. Carbamidomethylation of Cys was set as a fixed modification, with oxidation of Met set as a variable modification. The false-discovery rate (FDR) cutoff was 1% for all peptides.

**Immunoblot analysis**

For immunoblot analysis of fractions generated during LD isolation, percentage per total volume of each protein lysate fraction was collected as indicated in Figure 3B and precipitated in acetone overnight at -80°C. Washes, resuspensions, and heating were performed as outlined above. Proteins were separated on a homemade 4-15% SDS-PAGE gel and then transferred to a 0.45μm nitrocellulose membrane in Towbin SDS transfer buffer (25mM Tris, 192mM glycine, 20% methanol, 0.05% SDS; pH 8.2) using a Criterion tank blotter with plate electrodes (BioRad 1704070) set to 70V constant. Immediately following transfer, membranes were stained with PonceauS and cut using a clean razor blade. Membranes were blocked with 5% milk dissolved in TBS-T buffer, and primary antibodies were allowed to bind overnight at 4°C. Primary antibodies used for determining protein expression are as follows: Pma1 (Abcam ab4645; 1:1,000 dilution), Tubulin (Abcam ab6160; 1:10,000 dilution), Porin (ThermoFisher CAT 459500; 1:1,000 dilution), and Pet10/Plin1 (Joel Goodman’s lab, 1:1,000), also used in Gao, et al, JCB, 2017. Immunoblots were developed by binding HRP-conjugated anti-rabbit IgG (Sigma A0545; 1:10,000), anti-rat IgG (Abcam ab97057; 1:10,000), or anti-mouse IgG (Abcam ab6728; 1:2,000) secondary antibodies to the membrane for one hour in the presence of 5% milk followed by three washes in TBS-T and developing with ECL substrate (1705061 BioRad). Signal was captured by x-ray film.

**QUANTIFICATION AND STATISTICAL ANALYSIS**

**Statistical analysis**

All statistical analyses were performed using Graphpad Prism 8 software. Unpaired t-tests were performed with Welch’s correction. For one-way ANOVA, Brown-Forsyth and Welch ANOVA was performed followed by Turkey’s post-hoc test to extract p-values. For both t-tests and ANOVA, *p<0.05, **p<0.01, ***p<0.001. All graphs represent the mean ± standard deviation.

**Image quantification**

All image analysis was performed using Fiji (Schindelin et al., 2012). For Mander’s colocalization quantification, confocal microscopy images were split into respective channels and analyzed using the JACoP plugin (Bolte and Cordelières, 2006). M1 and M2 coefficient analysis was used with automatic thresholding. Relative M1 values were calculated with the average of log-phase M1 values as the reference.

**Proteomics quantification**

Proteomics quantification and analysis was performed using excel. All samples were analyzed in quadruplicate. To adjust for total protein differences between samples, the sum of all spectral counts within each sample was taken and divided by the average of the spectral count sums in log-phase LD fractions. This ensures differences observed in the proteomics data is not due to unequal ‘loading’ into the MS. To generate heat maps and volcano plots, Log2 values were calculated for the ratio of average protein expression in log-phase and AGR (i.e. Log2(protein A in AGR/protein A in log)). Confidence analysis on LD fractions was performed essentially as described previously (Bersuker and Olzmann, 2019). Specifically, the LD confidence score is a measurement of spectral counts for each protein in the LD fraction, subtracted by their spectral counts in the infranatant fraction. Therefore, the score reports on the most abundant proteins present in the LD fraction in a given growth condition. Confidence score is calculated as the product of two equations outlined below:

$$Equation \#1= \sum_{LD=1}^{LD=k} {(X}_{LD,P}-X_{Inf, P})$$

$$Equation \#2= \sum_{LD=1}^{LD=k} R_{LD,P}$$

In equation #1, X is the spectral abundance of a given protein in the LD fraction (X_LD,P_) or in the infranatant fraction (X_Inf, P_). Equation #2 describes the number of times a protein was detected in the LD fraction dataset for a given growth condition. For proteins that were more abundant in the infranatant fraction than the LD fraction, i.e. equation #1<0, values were manually set to zero. Difference in LD confidence scores was calculated by subtracting the AGR LD confidence score for each protein by the log-phase LD confidence score. Whole-cell abundance factors represent the sum of spectral counts for each protein in a given growth condition. Differences in whole-cell abundance factors represent protein abundance in AGR subtracted by protein abundance in log-phase.

**Cartoons development**

All cartoons created with BioRender.com.

**DATA AND CODE AVAILABILITY**

The subtomograms shown in Figure 1B-I were deposited in the Electron Microscopy Data Base (EMDB) with accession numbers EMD-24762, EMD-24764, EMD-24766, EMD-24767, EMD-24781, and EMD-24782, respectively.
